# Human breast milk extracellular vesicles from donors with asthma differentially the modulate release of inflammatory cytokines by primary human airway smooth muscle cells in a recipient-cell specific manner

**DOI:** 10.64898/2026.03.02.709065

**Authors:** Tamiris F. G. Souza, Taiana M. Pierdona, Samira Seif, Benjamin Bydak, Patience O. Obi, Joseph W. Gordon, Stuart Turvey, Elinor Simons, Piushkumar Mandhane, Theo Moraes, Padmaja Subbarao, Shreya A. Raghavan, Andrew J. Halayko, Meghan B. Azad, Ayesha Saleem

## Abstract

Breastfeeding provides health benefits in childhood, reducing the frequency of gastrointestinal and respiratory infections. Breastmilk (BM) is a rich source of bioactive molecules including extracellular vesicles (EVs), which exert immunomodulatory signalling in recipient cells, with cargo that is affected by maternal characteristics. Here we investigated the biophysical characteristics of BM-EVs from mothers with (asthmatic BM-EVs) or without asthma (control BM-EVs) and their effect on the release of cytokines from primary human hTERT-immortalized airway smooth muscle cells (hASMs) from asthmatic or non-asthmatic (control) donors. BM-EVs were isolated using size exclusion chromatography (N=5/group), characterized biophysically and by EV-specific protein markers. In addition, BM-EV were co-cultured (48h) with primary hASM cells from both non-asthmatic (control) and asthmatic donors to determine the effect on cytokine release. All participants were Caucasian and the BM was collected 12-15 weeks postpartum. BM-EVs showed the presence of intact and small-EVs (∼100 nm). Asthmatic BM-EVs appeared to have a smaller average EV size (135.6 nm) *vs.* controls (148.3 nm, p=0.0613), but ∼5-fold higher concentration of both total (p=0.0014) and small EVs (p=0.0016). The expression of EV subtype protein expression was reduced in asthmatic BM-EVs *vs.* control BM-EVs: CD63 by 86% (p=0.0224), flotillin-1 by 40% (p=0.0196), CD9 by 24% (p=0.0646) and HSP70 by 69% (p=0.0873). Asthmatic BM-EVs co-cultured with hASMs from control donors decreased pro-inflammatory cytokine release: MCP-1 by 55% (p=0.0286), IL-6 by 45% (p=0.0801) and IL-2 by 32% (p=0.0970) *vs.* control-BM-EVs. Conversely, asthmatic BM-EVs co-cultured with hASMs from asthmatic donors increased secretion of anti-inflammatory cytokine IL-10 by 32% (p=0.0660), and IL-1Ra by 75% (p=0.0875), and pro-inflammatory IL-2 by 57% (p=0.0688) *vs.* control-BM-EVs. Internalization of control and asthmatic BM-EVs was confirmed by labelled EV uptake experiments. No detrimental effects on cell viability with BM-EV treatment were observed. In summary, asthmatic BM-EVs are smaller and enriched in BM, and exert differential effects on cytokine release in a BM-donor and recipient-cell specific manner. Given that BM can enter infant airways, the immunomodulatory effects of BM-EVs on hASMs warrants further investigation to delineate the under underlying mechanisms.

## Introduction

Asthma is a substantial global health problem affecting more than 350 million children, adolescents, and adults worldwide (García-Marcos et al., 2023). It is a major pediatric non-communicable chronic disease that places a significant burden on financial and healthcare systems (Enilari & Sinha, 2019), with negative impacts on patient quality of life, such as a high loss of school days and requirements for urgent care (Serebrisky & Wiznia, 2019). Asthma is defined by a history of respiratory symptoms such as wheezing, shortness of breath, chest tightness and cough that vary over time and in intensity. Childhood asthma incidence and prevalence are high and may impair airway development, causing lung function deficits that persist into adulthood. However, morbidity and mortality are higher in adults (Dharmage et al., 2019). Some studies show that the natural history of asthma in almost 80% of cases begins during the first six years of life (Trivedi & Denton, 2019). Although asthma incidence in males is more prevalent in childhood, the pattern switches in adulthood with higher asthma cases reported in females (Dharmage et al., 2019). Early-life stimulants during fetal and infant development such as pollution, tobacco smoke, maternal health, nutrition, and lifestyle, can alter immune profiles increasing the risk for asthma and allergy in infants (Kozyrskyj et al., 2011; Subbarao et al., 2009). Breastfeeding appears to be protective for infant respiratory health, though the underlying mechanisms remain unclear (Miliku & Azad, 2018).

Breastfeeding provides immediate health benefits in early childhood and infancy, reducing the frequency of gastrointestinal and respiratory infections and hospitalizations (Christensen et al., 2020; Frank et al., 2019). Previous studies have reported protective effects of breastfeeding against the development and severity of asthma (Ahmadizar et al., 2017; Beasley et al., 2015; Dell & To, 2001; Dogaru et al., 2014; Hough et al., 2018; Klopp et al., 2017; Kull et al., 2004; Perikleous et al., 2022; Scholtens et al., 2009), with direct effects on lung growth and protection against wheezing in infants and in mothers with asthma (Azad, 2019; Azad et al., 2017; Dogaru et al., 2012). Conversely, some studies document no protection of breastfeeding against asthma and atopy (S. W. Burgess et al., 2006; Elliott et al., 2008; Kramer et al., 2007; Lossius et al., 2018; Sears et al., 2002). The molecular mechanisms underlying the protective effects of breastfeeding are still being understood.

Human breastmilk is an irreplaceable source of nutrition for optimal early infant growth. It contains macro- and micronutrients, and diverse bioactives including immune components, hormones, microbes, growth factors, oligosaccharides, and extracellular vesicles (Doare et al., 2018; Kim et al., 2020). Extracellular vesicles (EVs) are small messengers crucial for intercellular communication in physiological and pathological processes (Safdar et al., 2016; Tkach & Théry, 2016). EVs are delimited by a lipid bilayer and contain transmembrane and cytosolic proteins, lipids, and nucleic acids, such as miRNA, mRNA, and non-coding RNA (Colombo et al., 2014). All cells can release EVs with distinct cargo signatures as determined by donor cells, and exert functional effects in recipient cells (Esmaeili et al., 2022). Traditionally, EV classification was standardized by its biogenesis pathway: (1) exosomes (30-150 nm); or (2) ectosomes including (2a) microvesicles (150-1000 nm) and (2b) apoptotic bodies (1-5 μm) (Di Bella, 2022). Current international standardized guidelines for EV research recommend EVs to be categorized by size as small EVs (<200 nm) or large EVs (>200 nm), density, biochemical composition, and cellular origin (Welsh et al., 2024). The method of EV isolation and characterization can affect EV bio-composition which necessitates careful documentation of protocols, extensive characterization of EVs, and highlights the importance of using orthogonal approaches (Sidhom et al., 2020).

EVs have been found in every biological fluid tested (Couch et al., 2021), their small size facilitates translocation through physical barriers, and passage through the extracellular matrix (Gurunathan et al., 2021), allowing EVs to deliver encapsulated cargo at great distances without degradation (Kalra et al., 2016). The transfer of functional cargo via EVs between cells can influence several biological processes, such as cell motility, differentiation, proliferation, apoptosis, reprogramming, and immunity (Liu & Wang, 2023; Yáñez-Mó et al., 2015) in a donor- and recipient-cell dependent manner. Given that breastmilk EVs composition is affected by maternal characteristics (Kupsco et al., 2021), lactation period (Yun et al., 2021), maternal disease (Mirza et al., 2019), and stressors (Bozack et al., 2021), it is necessary to investigate if maternal asthma status can directly impact breastmilk EVs composition and downstream biological function. Human breastmilk EVs can be internalized by macrophages (Lässer et al., 2011) and non-transformed human intestinal epithelial crypt-like cells (HIEC) (Liao et al., 2017), and influence immune response (Admyre et al., 2007). The effects of EVs on T cell-mediated immune responses have been detected at both the protein (Yáñez-Mó et al., 2015; Zonneveld et al., 2021) and miRNA levels (Karlsson et al., 2016; Zhou et al., 2012). A recent study established that milk EVs can preserve bronchial epithelial cell integrity and reduce inflammatory cytokine release during a viral challenge (Karra et al., 2022). Quantitative proteomics has identified novel proteins involved in regulating cell growth and controlling inflammatory signaling pathways in human breastmilk EVs (Van Herwijnen et al., 2016). While the lion’s share of attention has been focused on elucidating the effects of breastmilk EVs in the gastrointestinal tract due to the oral ingestion of milk by infants, breastmilk EVs can also be delivered to human airways through inhalation of milk aerosolization by infant suckling and breathing patterns (Goldfield et al., 2006). It is therefore likely that breastmilk EVs can affect human airway smooth muscle (hASM) cells, which play an integral role in the pathogenesis of asthma by regulating airway remodelling and inflammation. Thus, here we sought to investigate changes in breastmilk EV composition as a function of asthma status in mothers, and evaluated putative immunomodulatory functional effects of these EVs on primary hASM cells obtained from donors with and without asthma.

## Materials and methods

### 1. Study participants and ethics statement

We obtained samples of breast milk (BM) at 13-15 weeks postpartum from mothers previously diagnosed with asthma (N=5) and those without asthma as controls (N=5), enrolled in the Canadian Healthy Infant Longitudinal Development (CHILD) Cohort Study, directed by site-lead Dr. Meghan Azad at the University of Manitoba (Azad et al., 2017; Moraes et al., 2015; Subbarao et al., 2015). The CHILD study is the largest longitudinal study in Canada exploring relationships between environment and genetics in the early-life development of allergies and asthma (Takaro et al., 2015). All women involved in the study gave their signed informed consent to participate, and were matched by age, BMI, race and age of milk (weeks) (**Table 1**). We excluded mothers with other chronic health conditions expected to influence EV production (e.g., diabetes, arthritis, and irritable bowel syndrome). The study was approved by the Human Research Ethics Board at the University of Manitoba to Dr. Ayesha Saleem (HS23677 (H2020:092)). Asthma in offspring was diagnosed by trained healthcare professionals (physicians, nurses, or clinical research associates at each site) based on a physical examination and detailed history of respiratory symptoms, such as wheezing, breathlessness, chest tightness and cough (Miliku & Azad, 2018). Maternal age, ethnicity, education, prenatal smoking, BMI, diabetes (type 1, type 2, or gestational), parity (number of previous live births), and asthma status were self-reported during pregnancy.

**Table 1.** Characteristics of the subjects.

| **Characteristics** | **Control**<br>**(N=5)** | **Asthma**<br>**(N=5)** | **Participants**<br>**(N=10)** |
| --- | --- | --- | --- |
| Age (years) | 31.2<br>(29 – 32) | 32.8<br>(25 - 36) | 32<br>(25 – 36) |
| BMI (kg/m<sup>2</sup>) | 23.3 | 23.1 | 23.2 |
| Race (%) |  |  |  |
| Caucasian | 100 | 100 | 100 |
| Food allergy (%) | 0 | 80 | 40% |
| Weeks milk age (weeks) | 13.5<br>(12.7 – 14.6) | 13.9<br>(12.9 – 15.6) | 13.72<br>(12.7 – 15.6) |

### 2. Collection and processing of breastmilk samples

A volume of 10 ml of breastmilk from participants was collected at the home visit (one sample per mother) 13-15 weeks post-partum (range of 12.7 to 15.6 weeks), divided into aliquots, frozen at −80 °C and then stored in liquid nitrogen (Moraes et al., 2015). Frozen milk samples were thawed and prepared for EV isolation as follows. Milk samples were centrifuged at 2,000x *g* for 10 min at 4 ^◦^C to remove fat globules (16K Microcentrifuge, Bio-Rad Laboratories). Next, the supernatant was centrifuged at 12,000x *g* for 30 min (4 ^°^C), passed through a 0.22 μm filter to eliminate possible cellular debris and contaminants and stored at −80 ^°^C until EV isolation (Bickmore & Miklavcic, 2020). The workflow for breastmilk EV (BM-EV) isolation and characterization is shown in **Fig. 1A**.

**Figure 1.**
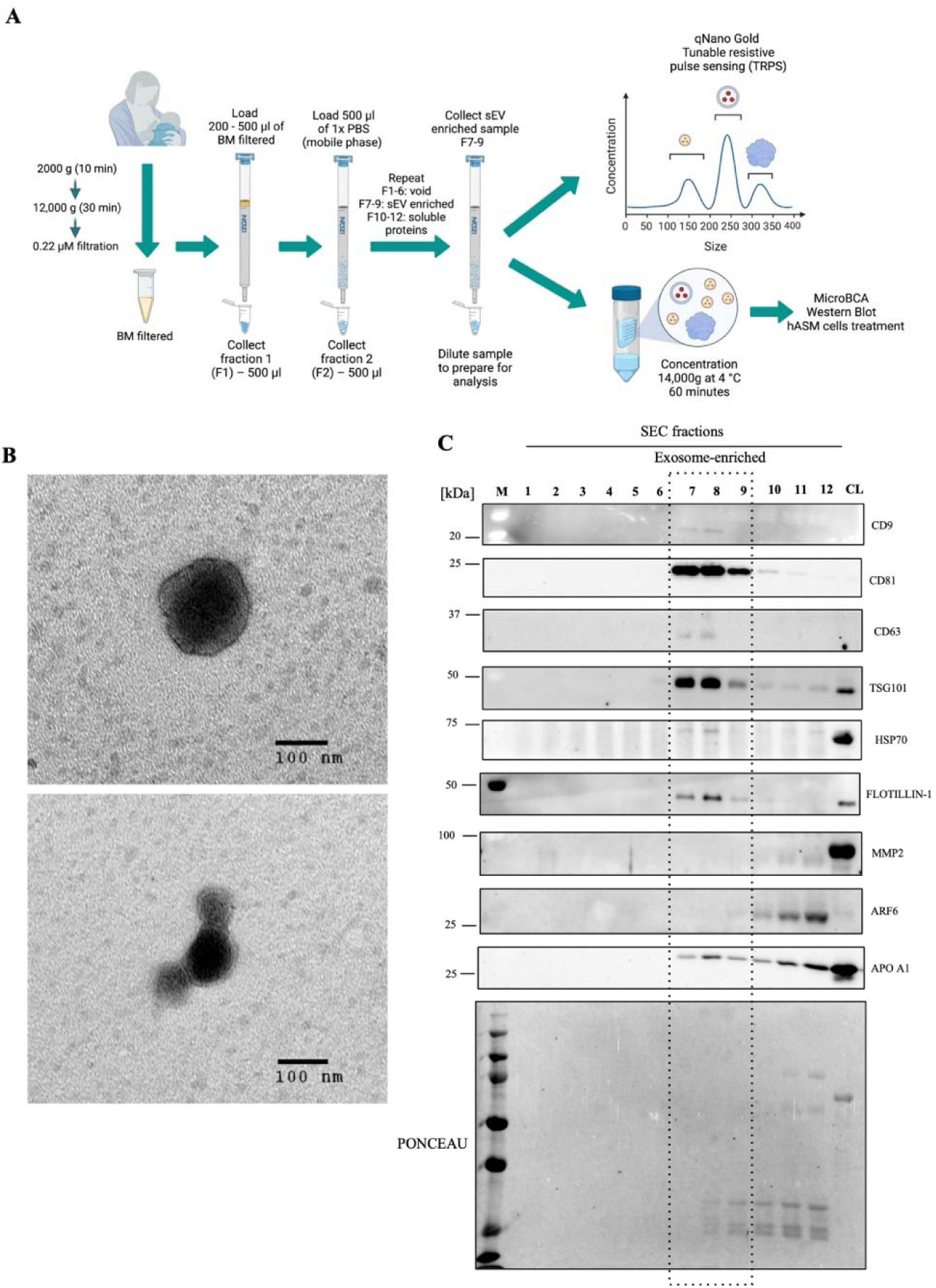
Breast-milk extracellular vesicles (BM-EVs) isolation, TEM imaging and expression of general EV marker proteins in EV fractions. (A) Schematic representation of BM-EVs isolation and characterization. BM was collected from 3-4 months post-partum mothers with and without asthma diagnosis. BM-EVs were isolated by SEC (qEV original columns, Izon Science Ltd) and analyzed for size, concentration and zeta potential through TRPS using the qNano Gold instrument (Izon Science Ltd.). BM-EVs were concentrated for protein yield quantification through MicroBCA Protein Assay kit (Thermo Scientific), Western blot analysis for exosome and microvesicles markers and hASM cells treatment for cytokines release analysis. (B) TEM showing exosomal morphology and approximate size of BM-EVs (scale bar 100[nm). (C) Western blotting was performed (12% SDS-PAGE) and Ponceau S staining used as a loading control. Proteins enriched in exosomes (CD9, CD81, CD63, TSG101, HSP70 and Flotillin-1), in microvesicles (MMP2 and ARF6), and lipoproteins (APO-A1) were analysed. F7 to 9 were considered exosome-rich while microvesicles and lipoprotein-poor. CL: cell lysate from breast tissue, M: marker lane.

### 3. BM-EV isolation and characterization

BM-EVs were isolated using size exclusion chromatography (SEC) (qEV original columns, Izon Science Ltd) according to the manufacturer’s instructions. Briefly, 500 μl of the filtered breastmilk was loaded onto qEV columns and fractions (F) were collected with filtered phosphate-buffered saline (PBS) as the elution buffer (mobile phase). Each fraction collected (F1–12) was analyzed for protein yield and presence of EV specific markers as well as negative controls by Western blotting. The pooled fractions F7-9 were determined to be enriched with small-EVs and used for subsequent functional analyses.

#### 3.1. Western blotting

BM-EV preparations (F1-12) were denatured by the addition of β-mercaptoethanol and 4x laemmli buffer, followed by incubation at 95 ^°^C for 5 min. 5 μg of total protein was loaded on 12% SDS-PAGE gels and electrophoresed at 90 mV for 30 min and then 120 mV for 1.5 h. Proteins were transferred to a nitrocellulose membrane using Trans-Blot® Turbo™ (Bio-Rad Laboratories). Next, the membrane was incubated in Ponceau S staining for 5 min, and a colorimetric imaged taken using the ChemiDoc^TM^ MP Imaging System (Bio-Rad Laboratories) to serve as loading control. Ponceau staining was removed by washing the membrane for 1 min in ultrapure water. Membrane was then blocked for 1 h at room temperature using 5% skim milk in Tris-buffered saline Tween-20 solution (TBST), and incubated with primary antibodies in 1% skim milk overnight, at 4 °C against target proteins to evaluate the purity and expression of sEV preparations. The following primary antibodies were used: rabbit polyclonal anti-TSG101 (T5701, Sigma-Aldrich Co, 1:200), rabbit polyclonal anti-CD63 (SAB4301607, Sigma-Aldrich Co, 1:200), mouse monoclonal anti-CD81 (sc-166029, Santa Cruz Biotechnology, 1:200), mouse monoclonal anti-CD9 (CBL162, Sigma-Aldrich Co, 1:100), rabbit polyclonal anti-Flotillin-1 (F1180, Sigma-Aldrich Co, 1:200), mouse monoclonal anti-heat shock protein 70 (HSP70) (H5147, Sigma-Aldrich Co, 1:500), mouse monoclonal anti-ARF6 (sc-7971, Santa Cruz Biotechnology, 1:200), mouse monoclonal anti-MMP2 (sc-13595, Santa Cruz Biotechnology, 1:200) and mouse monoclonal anti-Apolipoprotein A1 (Apo-A1) (0650-0050, Bio-Rad Laboratories, 1:200). Membranes were washed 3 times with TBST and incubated with anti-mouse IgG secondary antibody (A16017, Thermofisher, 1:10,000) or anti-rabbit IgG secondary antibody (A16035, Thermofisher, 1:1,000) for 1 h at room temperature in 1% milk. Clarity™ ECL substrate (Bio-Rad Laboratories) solutions were applied to visualize bands via enhanced chemiluminescence, and images captured using the ChemiDoc^TM^ MP Imaging System (Bio-Rad Laboratories). Band intensity was quantified using Image lab software (Bio-Rad Laboratories) and corrected for loading using Ponceau S. The pooled fractions F7–9 were considered small EV (sEV)-rich, and lipoprotein-poor, and were pooled to characterize EVs according to MISEV guidelines (Welsh et al., 2024): by size, concentration and stability (zeta potential) using characterized through Tunable Resistive Pulse Sensing, and size and morphology by transmission electron microscopy.

#### 3.2. Transmission electron microscopy (TEM)

10 µl of freshly isolated BM-EVs (F7-9) were fixed by mixing aliquot with an equal volume of 4% paraformaldehyde in PBS. Then, 5 μl of the BM-EVs suspension was added to a clean parafilm. Formvar/Carbon, grids (Cat# 01800-F, Electron Microscopy Sciences) were floated with their coated side facing the sample suspension and incubated for 20 min at room temperature. Grids were washed with 100 μl of PBS for 30 s where PBS was added to a parafilm, and grids transferred with forceps onto PBS droplets with the membrane side down. Next, 50 μl of 1% glutaraldehyde in PBS was added to the grids and incubated for 5 min. To remove the excess fixation buffer, the grids were washed 8 times with 100 μl of distilled water as described above. Grids were then stained with 1% uranyl acetate for 2 min and left to dry at room temperature. Grids were observed under the electron microscope at 60 kV and images were taken at 64000x magnification using Phillips CM10 Electron Microscope.

#### 3.3. Tunable Resistive Pulse Sensing (TRPS)

The size, concentration, and zeta potential of BM-EVs were measured using a TRPS system (qNano Gold, Izon Science Ltd., New Zealand). TRPS is based on the detection of an electrical resistive pulse caused by individual nanoparticles passing through a nanopore, which is detected and measured by the instrument. The magnitude and number of the pulses or blockades are converted to particle diameter and concentration, respectively (Vogel et al., 2017). Zeta potential is an indicator of the colloidal stability of particles in a dispersed system through the measurement of electrically charged particles. The electrophoretic mobility in a suspension can be measured and indicates the stability of particle-particle and particle-medium interactions (Midekessa et al., 2020). The carboxylated polystyrene particles (CPC) with known average size and concentration were used to calibrate the data acquired from the particle suspension (Vogel et al., 2016). An aliquot of 35 μl of each BM-EV suspended in electrolyte solution was used to characterize nanoparticles using an NP150 nanopore and 47.0 stretch, using a combination of different pressures and a constant voltage. Data was collected and analyzed by the Izon Science Control Suite v. 3.3.

#### 3.4. Protein quantification

Breastmilk sEV preparations (F7-9, roughly 1.5 ml) were loaded onto Amicon Ultra-4 3 kDa centrifugal filters (Merck Millipore), and centrifuged at 14,000x *g*, for 60 min at 4 ^°^C to concentrate EV isolates (∼ 50 µl) for further assays (Pierdoná et al., 2022). sEV preparations were lysed and total proteins extracted using 1:1 Pierce RIPA solution with protease inhibitor tablet (Roche). Samples were vortexed for 5s, followed by two freeze/thaw cycles with liquid nitrogen, sonicated at 30 Hz 3×3s on ice and centrifuged at 21,130x *g* for 20 min at 4 ^°^C. Protein concentration was measured using the Pierce™ Micro BCA Protein Assay kit (Thermo Scientific), in accordance with the manufacturer’s instructions. Briefly, the working reagent was prepared by mixing 25 parts of Reagent A, 24 parts of Reagent B and 1 part of Reagent C. The standards were prepared by serial dilution of 2 mg/ml bovine serum albumin (BSA) ampule into clean vials using ultrapure water. 150 µL of each standard or sample was added to a 96-well microplate in duplicates, followed by the addition of 150 µL of working reagent to each well. The samples were incubated 37 ^°^C for 2 h and absorbance was measured at 562 nm using a microplate spectrophotometer (BioTek Epoch). Total EV protein yield (µg) was determined by multiplying the protein concentration (µg/µl) with the volume (µl) of the purified EV preparations. EV protein yield (pg/particle) was obtained by dividing the total protein yield by the total number of particles for each sample.

### 4. BM-EV treatment in human airway smooth muscle (hASM) cell culture from asthmatic and non-asthmatic (control) donors

Primary hASM cells from asthmatic and non-asthmatic controls were generously donated by Dr. Andrew Halayko (University of Manitoba). These cells were isolated from lung tissue obtained through lung resection surgery following approval from the Human Research Ethics Board of University of Manitoba. Cells from two donors were used, one with no previous history of chronic lung disease (asthma/COPD) designated as control (hASM control) and one diagnosed with asthma (hASM asthma). G418 was added to the cells to induce immortalization through stable ectopic expression of human telomerase reverse transcriptase (hTERT) (J. K. Burgess et al., 2018). To induce phenotype differentiation that partially resembles hASM cells *in vivo*, 24 h after seeding the cells, the culture medium was replaced by serum starved DMEM supplemented with 1% insulin-transferrin-selenium (ITS) for 7 days (Pascoe et al., 2021). The suspension of hASM cells were seeded at a concentration of 2×10^4^ cells/ml into 6-well (multiplex assay) and 24-well (MTT assay) plates in DMEM containing 10% fetal bovine serum (FBS, Gibco), 1% penicillin-streptomycin (P/S, Gibco) and 400 μg/ml of G418 (Sigma). Cells were used between passages 5-8. For a spike in experiments, both control and asthma hASM cells were co-cultured for 48 h with BM-EVs. An aliquot of 200 μl of filtered BM was diluted in 300 μl of PBS and BM-EVs were isolated as described above. BM-EVs from both mothers with asthma (N=5) and without asthma (N=5) were used to treat hASM cells (from control and asthmatic donors). The treatment with BM-EVs was done in the presence or absence of interleukin 1-beta (IL-1β, 10 ng/ml) for 24 h, as a challenge to stimulate pro-inflammatory cytokine release.

#### 4.1. Cell viability

Cell viability was evaluated by using the 3-(4,5-dimethyl thiazolyl)-2,5-diphenyltetrazolium bromide (MTT) assay (Mosmann, 1983). After 48 h of BM-EVs treatment, 300 μl of MTT solution (5 mg/mL) was added to the hASM cells and incubated at 37 ^°^C for 3 h. Then, medium containing MTT was removed, and 300 μl of DMSO was added. Absorbance was determined using a microplate reader (BioTek Epoch) at a wavelength of 540 nm. The results were calculated by the absorbance of treated cells over the control cells (vehicle, PBS) x 100%.

#### 4.2. Cytokine analysis using multiplex analysis

The conditioned media from hASM control and asthma cells after BM-EV treatment was collected to quantify inflammatory cytokines release of the following: granulocyte-macrophage colony-stimulating factor (GM-CSF), interferon-gamma (IFN-γ), IL-1β, IL-1receptor agonist (IL-1Ra), IL-2, IL-4, IL-5, IL-6, IL-8, IL-10, IL-12p40, IL-12p70, IL-13, monocyte chemoattractant protein-1 (MCP-1), tumor necrosis factor-alpha TNFα). Cells were treated with BM-EVs (from mothers with and without asthma) for 48 h, with and without pre-incubation with IL-1β for 24 h, prior to conditioned media collection. Cytokine levels (pg/ml) were analyzed using the multiplex Human Focused 15-Plex Discovery Assay (Eve Technologies, Alberta, Canada) (Pascoe et al., 2021).

### 5. BM-EV uptake by hASM cells

To confirm BM-EV uptake by hASM cells, BM-EVs from mother with and without asthma (control) were labelled using PKH67 lipophilic membrane dye kit (Sigma-Aldrich Co) as before (Lässer et al., 2011). PKH67 dye has been successfully used to label exosomes and EVs for *in vitro* and *in vivo* tracking experiments (Takov et al., 2017). Briefly, BM-EV isolates (40 µl) were volumed up to 500 µl with Diluent C (EV solution). Then, 2 µl of PKH67 dye (1 µM) was added to the 498 µl of the diluted EV mix, mixed immediately by gentle pipetting for 30 s and incubated at room temperature for 5 min. The staining process was stopped by adding 500 µl of FBS and incubated for 1 min to quench excess dye. Next, 500 µl of DMEM (FBS and P/S free) was added and BM-EVs re-isolated by differential ultra-centrifugation (dUC, 70 min at 100,000 x *g*, Sorvall MTX 150 Micro-Ultracentrifuge, Thermo Scientific using a fixed angle rotor, S110-A, Thermo Scientific™ Sorvall™ MTX) to obtain labelled BM-EVs and remove non-specific dye aggregates. The pellet containing labelled BM-EVs was resuspended using 50 µl of PBS, and labelled EVs added to 10K hASM cells in chamber slide, (Cat # 154526, Thermo Scientific™) for 24 h. After the incubation, cells were washed twice with PBS and counterstained with Hoechst for 30 min in the dark. In the last step, cells were washed twice with PBS and 1 ml of DMEM added to cells in each chamber and imaged using a Zeiss LSM700 spectral confocal microscope, at 20X magnification.

### 6. Statistical analyses

Average EV size, zeta potential, EV concentration, protein yield and expression of EV markers were analyzed using an unpaired t-test. Two-way ANOVA was used to analyze data from cell viability assay. Cytokine levels were analyzed in the following manner: data were tested for normality using a Shapiro-Wilk test. Data normally distributed were analyzed by unpaired Student’s t-test, followed by Welch’s correction. For data non-normally distributed, the non-parametric Mann-Whitney test was used. All data were analyzed using PRISM software, version 9 (GraphPad, San Diego, CA) with 95% confidence intervals. Significance was set at p<0.05 and data are expressed as mean ± standard error (N=4-5).

## Results

### Characteristics of participants

All participants (N=10) were self-reported Caucasian, with an average age of 32 years (range of 25 to 36) and an average BMI of 23.2. The lactational stage was between 12 and 15 weeks (average of 13.7 weeks). 80% of mothers with asthma also reported presence of food allergy (**Table 1**).

### Morphology of BM-EVs, purity and characterization of small EV-rich SEC fractions

EVs were isolated and characterized according to MISEV guidelines (Welsh et al., 2024) and as shown in **Fig. 1A**. The morphology of BM-EVs as measured by TEM (**Fig. 1B**), demonstrates isolation of intact EVs with a distinct outer membrane and spherical-shape, with approximate diameter of 100 nm (**Fig. 1B**). EV subtype marker expression was analyzed by Western blotting to validate SEC-EVs isolation from Fractions (F1-12, **Fig. 1C**). Equal amounts of total BM-EVs proteins were resolved on a 12% SDS-PAGE and probed for expression of proteins enriched in exosomes (CD9, CD81, CD63, TSG101, HSP70 and Flotillin-1), microvesicles (MMP2 and ARF6) and lipoproteins (ApoA1) markers. F7-9 were considered exosome or small-EV-rich due to the presence of CD-9, CD81, TSG101, Flotillin-1 and HSP70, microvesicles- and lipoprotein-poor. Total protein yield from each fraction (F1-12) showed an exponential increase from F7 onwards in both groups in line with previous reports (Boing et al., 2014, **Fig. S1**). SEC F10-12 presented higher protein content but did not show the expression of small EV-specific markers, likely due to increased protein contamination in EV preparations as reported before (Vogel et al. 2016) (**Fig. 1C**, and **Fig. S1**).

### Biophysical characteristics of BM-EVs isolated from donors with or without asthma

Average EV size appeared to decrease in BM-EVs from asthmatic donors (135.6 ± 1.32 nm) *vs*. control-EVs (148.3 ± 8.02 nm, p=0.0613, **Fig. 2A**). Zeta potential indicative of stability of nanoparticles in a suspension did not show difference between control (−12.46 ± 0.41 mV) and asthma BM-EVs (−12.76 ± 0.13 mV, **Fig. 2B**). Total EV concentration was ∼5-fold higher in asthmatic BM-EVs (2.39×10^11^ particles/ml) compared to controls (5.13×10^10^ particles/ml, p=0.0014, **Fig. 2C**). EV size distribution and concentration histogram showed a shift to the left in asthmatic BM-EVs between 100–200 nm range compared to control BM-EVs (**Fig. 2D**). In line with this, the concentration of small BM-EVs (0-200nm) was ∼5-fold higher in asthma (2.29×10^11^ particles/ml) compared to control (4.41×10^10^ particles/ml, p=0.0016, **Fig. 2E**), while large BM-EVs was similar between groups (p=0.3012, **Fig. 2F**). Pooled BM-EV-rich fractions (F7-9) from asthma and control groups were concentrated by centrifugation and characterized by total protein content. Total BM-EV (F7-9) protein yield was similar between asthma (29.2 ± 2.3µg) and control (39.4 ± 5.9µg, p=0.2717, **Fig. 2G**), and remained unchanged when corrected by the number of EVs (**Fig. 2H**). We conclude that while there is an increase in BM-EV concentration with asthma, the protein content of BM-EVs per vesicle remains unaltered.

**Figure 2.**
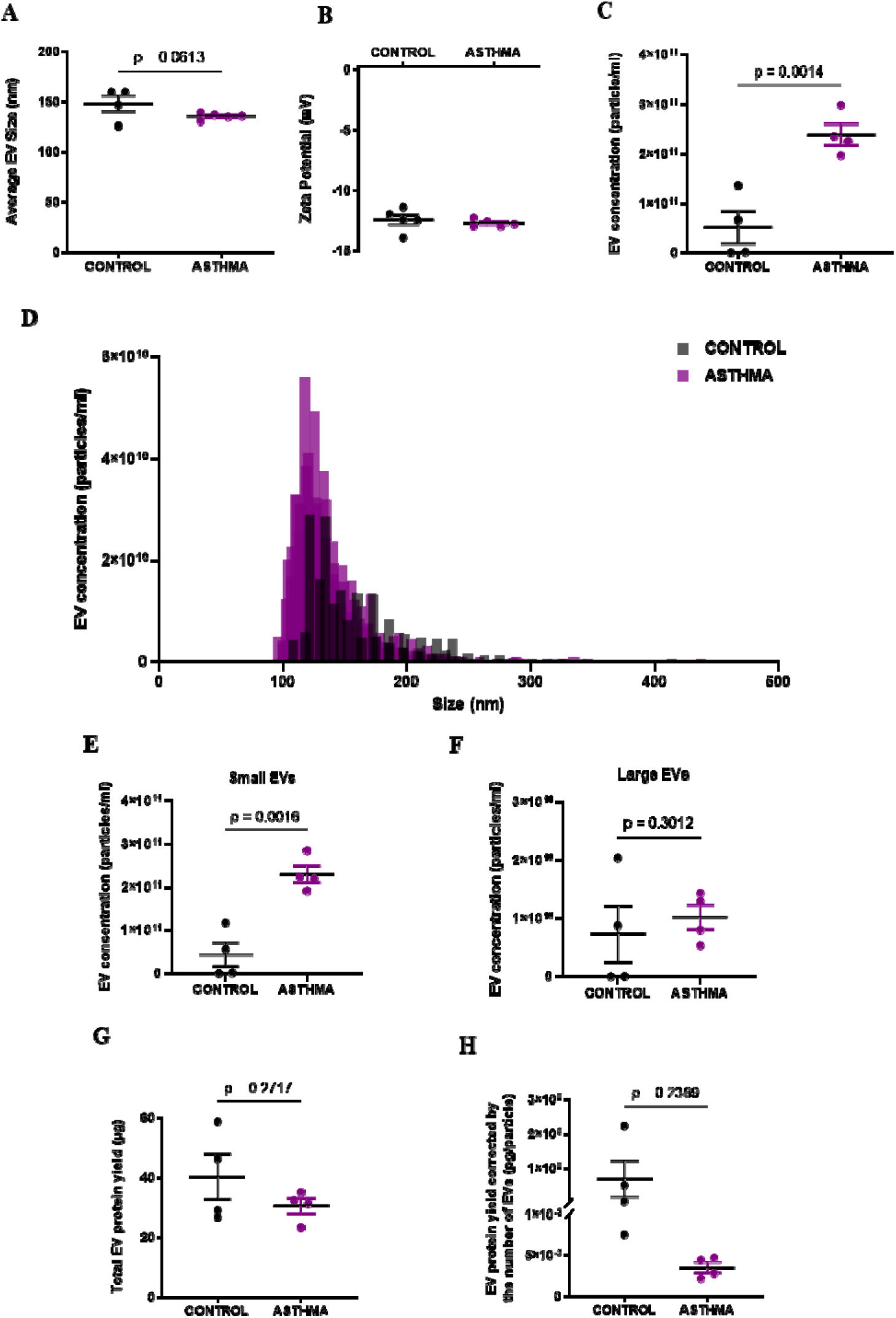
Control and asthma BM-EVs biophysical characterization by size, zeta potential, concentration, and protein content. **(A)** Average EV size was smaller in BM-EVs from mothers with asthma (asthma BM-EVs) vs. mothers without asthma (control BM-EVs) (p=0.0613). **(B)** Zeta potential was unchanged between groups. **(C)** The concentration of asthma BM-EVs was ∼5-fold higher (p=0.0014) compared to the control group. **(D)** EV size distribution (0-500 nm) of particles by concentration (particles/ml) demonstrates asthma BM-EVs are enriched with smaller EVs (100-150nm) compared to control BM-EVs. **(E)** Total concentration of small EVs (0-200 nm) was ∼5-fold higher in asthma BM-EVs compared to control group (p=0.0016). **(F)** Total concentration of large EVs (200-500 nm) was similar between groups (p=0.3012). **(G)** The total EV protein yield (µg) of the combined fractions F7-9 was unchanged between groups. **(H)** Similarly, the protein content corrected by the number of particles of BM-EVs was comparable between groups (pg/particle). Data were analyzed using an unpaired Student’s t-test and expressed as mean ± standard error (N=4-5).

### EV subtype-specific expression of marker proteins in BM-EVs from mothers with or without asthma

Next, EV specific and non-specific protein expression was analyzed by Western blotting using asthma and control BM-EV (F7-9) protein lysates. Equal amounts of proteins from each were resolved using 12% SDS-PAGE and ponceau or Coomassie staining was used as a loading control. Representative immunoblots (**Fig. 3A**) and graphical representation of quantified results (**Fig. 3B-H**) is included. Asthmatic BM-EVs had decreased levels of small EV protein markers including CD63 by ∼86% (p=0.0224, **Fig. 3C**), CD9 by ∼24% (p=0.0646, **Fig. 3D**), flotillin-1 by ∼40% (p=0.0196, **Fig. 3E**), and HSP70 by ∼69% lower (p=0.0873, **Fig. 3G**) compared to control BM-EVs. To evaluate the presence of microvesicle-specific proteins, we measured MMP-2 expression, but were unable to detect in our EV lysates (**Fig. 3A**). There was no difference between groups for other small EV markers such as CD81 (**Fig. 3B**) and TSG101 (**Fig. 3F**), and lipoprotein-marker ApoA1 (**Fig. 3H**). ApoA1 expression is used as a proxy measure to illustrate the purity of EV isolation by SEC, detecting the presence of lipoproteins co-isolated with BM-EVs (**Fig. 3H**).

**Figure 3.**
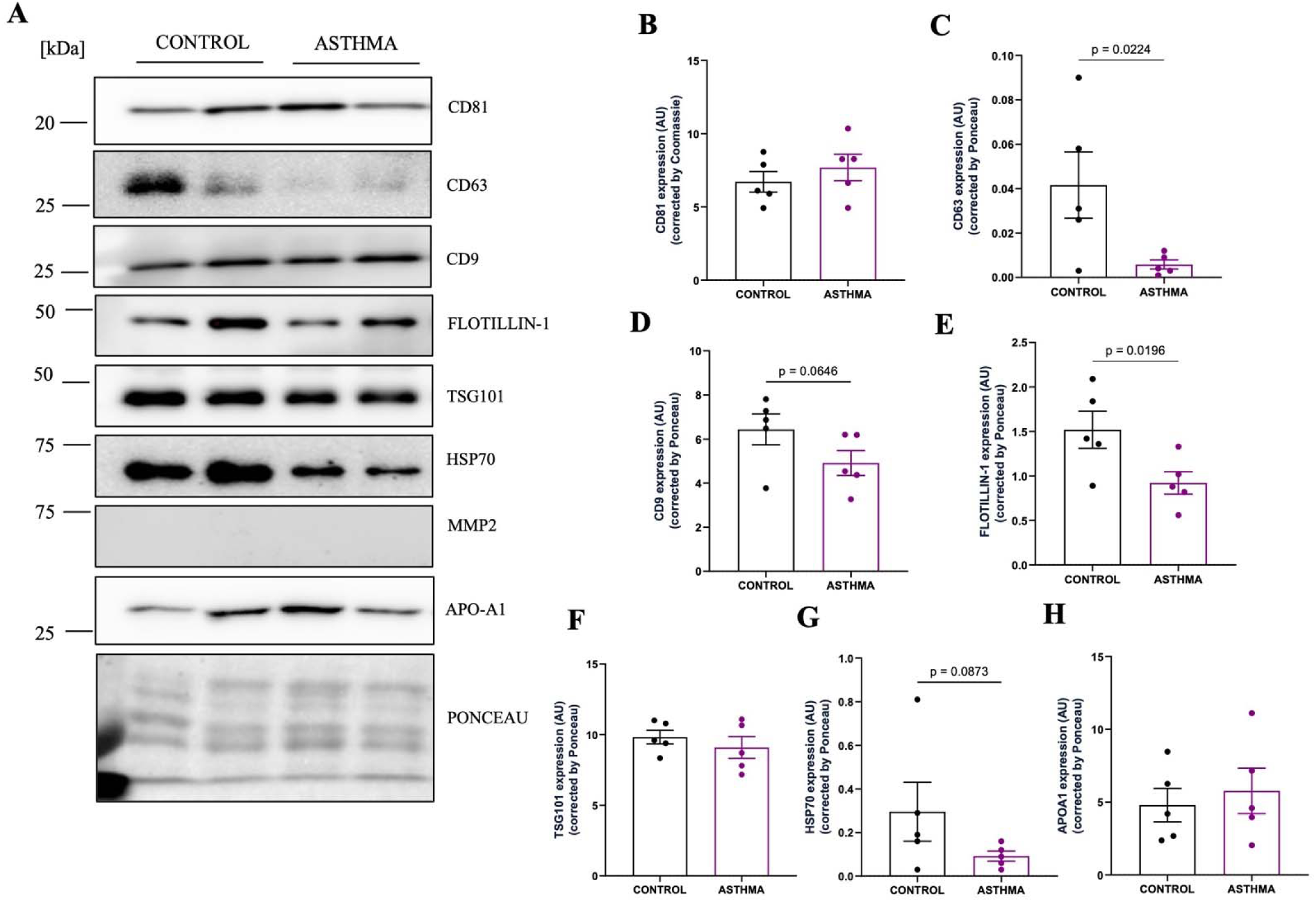
Expression of specific and non-specific EV subtype markers in BM-EVs from mothers with and without asthma. **(A)** Equal amounts of protein (5 µg/mL) of BM-EVs from mothers with and without asthma were subjected to SDS-PAGE (12% SDS) and the expression of proteins enriched in *exosomes* [CD81 (22 kDa), CD63 (28 kDa), CD9 (25 kDa), Flotillin-1 (48 kDa), TSG101 (46 kDa), HSP70 (70 kDa)], in *microvesicles* [MMP2 (63kDa)], and in *lipoproteins* [APO-A1 (25kDa)] were quantified. Ponceau or Coomassie staining was used as a loading control to normalize protein content for all markers. **(B)** CD81 levels remained unchanged between the groups, **(C)** while asthma BM-EVs expressed ∼86% lower CD63 (p=0.0224) and **(D)** ∼ 24% lower CD9 (p=0.0646). **(E)** Flotillin-1 expression was lower by ∼40% in asthma BM-EVs compared to control (p=0.0196). **(F)** TSG101 levels remained unchanged, **(G)** while HSP70 was ∼69% lower in the asthma BM-EV (p=0.08). **(H)** APO-A1 was used as a purity control to investigate the presence of lipoproteins in the samples and did not show difference between groups. BM-EVs did not show any expression for MMP2 protein. Data were analyzed using an unpaired Student’s t-test with p<0.05 considered as significant and expressed as mean ± standard error (N=5).

### Cell viability of hASM cells co-cultured with BM-EVs

hTERT-modified primary human airway smooth muscle (hASM) cells isolated from lung tissue from asthmatic and non-asthmatic (control) donors were obtained from Dr. Andrew Halayko’s repository at the University of Manitoba. Primary asthmatic and control hASM cells were co-cultured with BM-EVs isolated from asthmatic and control donors for 48 h, and cell viability measured by the MTT assay. PBS was included as a negative control, and Triton-X 0.1% as a positive control. The percent viability of cells treated with control and asthma BM-EVs remained unchanged in control hASM cells (93.61 ± 3.8%; 102.7 ± 4.39%) and asthmatic hASM cells (95.08 ± 2.39%; 99.97 ± 3.08%, **Fig. 4**), respectively. Treatment with 0.1% Triton X-100 showed significant reduction in cell viability in both control and asthmatic hASM cells groups to 8.2 ± 0.1% and 9.7 ± 0.6%, respectively, when compared with negative control (PBS). To confirm uptake of BM-EVs by hASM cells, control and asthmatic BM-EVs were labelled with PKH67 and were co-cultured with hASM cells for 24 h. Cells were subsequently stained with Hoechst and imaged on a confocal microscope. Uptake of control and asthmatic BM-EVs is shown in representative fluorescent images (**Fig. S2**).

**Figure 4.**
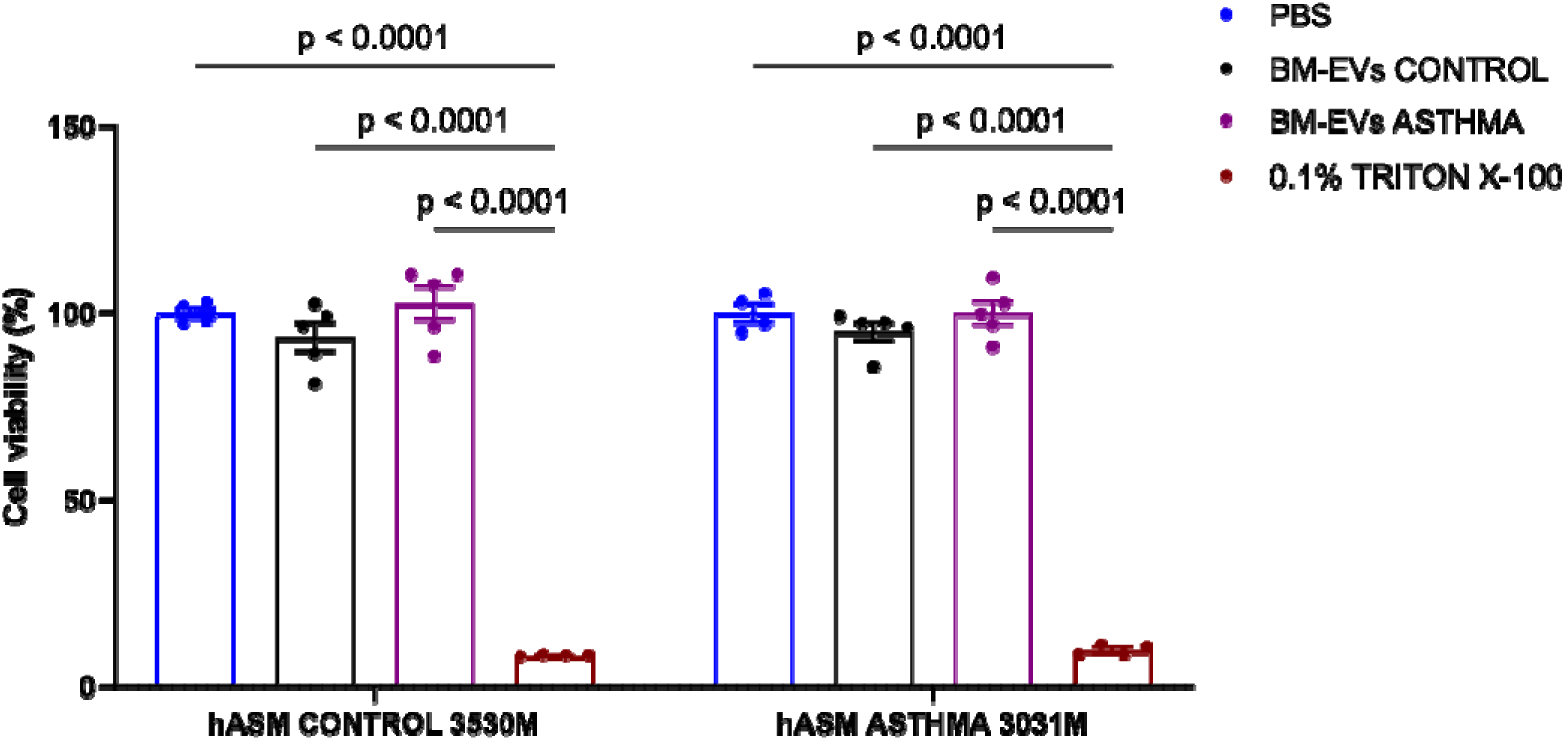
Cell viability of hASM cells treated with BM-EVs. MTT assay was performed of hASM cells from an asthmatic and non-asthmatic (control) donors and treated with BM-EVs from mothers with and without asthma (control). The cell viability (%) of hASM cells was expressed after 48h of incubation with BM-EVs, PBS (negative control), or 0.1% Triton X-100 (positive control). Control and asthma BM-EVs treatment did not affect the cell viability of hASM cells, and only the positive control reduced by ∼90%. Data were analyzed using two-way ANOVA and expressed as mean ± standard error (N=4-5). hASM = human airway smooth muscle.

### Cytokine release by hASM cells treated with control and asthma BM-EVs

Conditioned media from control and asthmatic hASM cells was collected to investigate the effects of BM-EV treatment on the release of chemokines and cytokines after treatment for 48 h. GM-CSF, IFN-γ, IL-1β, IL-1Ra, IL-2, IL-4, IL-5, IL-6, IL-8, IL-10, IL-12p40, IL-12p70, IL-13, MCP-1 and TNFα released by hASM cells with or without IL-1β exposure were measured by multiplex assay. All cytokine measurements are available in **Table 2**. Asthmatic BM-EVs co-cultured with hASMs from control donors reduced pro-inflammatory cytokine release: MCP-1 by 55% (p=0.0286, **Fig. 5A**), IL-2 by 32% (p=0.0970, **Fig. 5C**) and IL-6 by 45% (p=0.0801, **Fig. 5D**) *vs*. control BM-EVs. Conversely, asthmatic BM-EVs co-cultured with hASMs from asthmatic donors increased anti-inflammatory cytokine release: IL-1Ra by 75% (p=0.0875, **Fig. 5B**) and IL-10 by 32% (p=0.0660, **Fig. 5F**) and pro-inflammatory IL-2 by 57% (p=0.0688, **Fig. 5C**) *vs*. control BM-EVs. BM-EV treatment did not affect IL-8 (**Fig. 5E**), nor GM-CSF levels in either cell type (**Fig. 5G**). To trigger the release of chemokines and cytokines by hASM cells and simulate airway inflammation during asthma, IL-1β was added 24 h before conditioned media collection (purple inset in Fig. 5). Asthmatic BM-EVs appeared to increase IL-4 release by 21% (p=0.0992, **Fig. 5H**) in control hASM cells only, while IL-6 levels did not change with either treatment or cell type (**Fig. 5I**).

**Figure 5.**
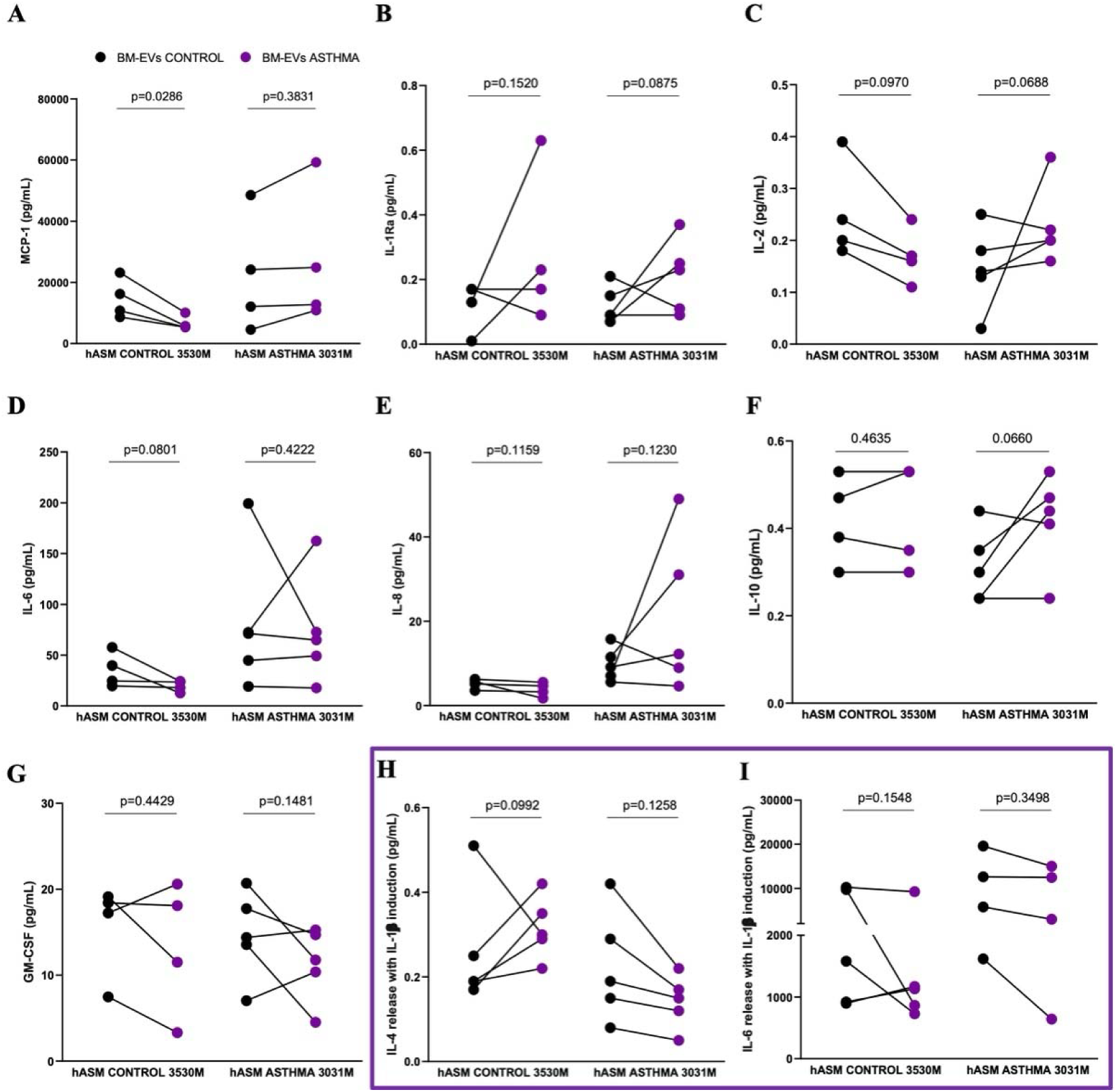
BM-EVs from donors with/without asthma modulated the release of inflammatory cytokines from primary hTERT-modified hASMs in a donor- and recipient-cell specific manner. Human airway smooth muscle (hASM) cells from two donors (control 3530M; asthma 3031M) were cultivated to investigate cytokine levels after co-culture with BM-EVs from mothers with or without asthma. Conditioned media was collected after 48h of incubation with BM-EVs, and cytokine levels (pg/ml) were analyzed using a multiplex assay (Eve Technologies). (**A**) Asthmatic-donor BM-EV treatment reduced MCP-1 release by 55% in non-asthmatic hASMs *vs*. control BM-EVs (p=0.0286). (**B**) However, asthmatic-donor BM-EVs showed an increase in IL-1Ra by 75% (p=0.0875) and (**C**) in IL-2 by 57% (p=0.0688) in asthmatic hASMs, and (**D**) a reduction in IL-6 by 45% in non-asthmatic hASMs *vs.* control BM-EVs (p=0.0801). (**E**) BM-EVs treatment did not affect IL-8 levels in both hASMs conditions. (**F**) Finally, asthmatic-donor BM-EVs increased IL-10 level by 32% (p=0.0660) in asthmatic hASMs, (**G**) while GM-CSF levels remained unaltered irrespective of treatment or cell type. (**H-I**) To trigger the release of chemokines and cytokines by hASMs and simulate airway inflammation during asthma, IL-1β was added 24h before conditioned media collection (purple box). (**H**) Asthma BM-EVs showed a slight increase in IL-4 (p=0.0992) in non-asthmatic hASM cells, (**I**) while IL-6 levels did not change with either treatment nor cell type. Overall, treatment with asthmatic donor-derived BM-EVs reduced pro-inflammatory cytokines (MCP-1, IL-6 and IL-2) in non-asthmatic hASMs, and increased anti-inflammatory cytokine (IL-10, IL-1Ra) and pro-inflammatory IL-2 in asthmatic-hASMs. Data normally distributed was analyzed by unpaired t-test, followed by Welch’s correction. For data non-normally distributed, the non-parametric Mann-Whitney test was used. Data were expressed as individual points for non-asthmatic hASM (3530M) and asthmatic hASM (3031M) cells (N=4-5).

**Table 2.**
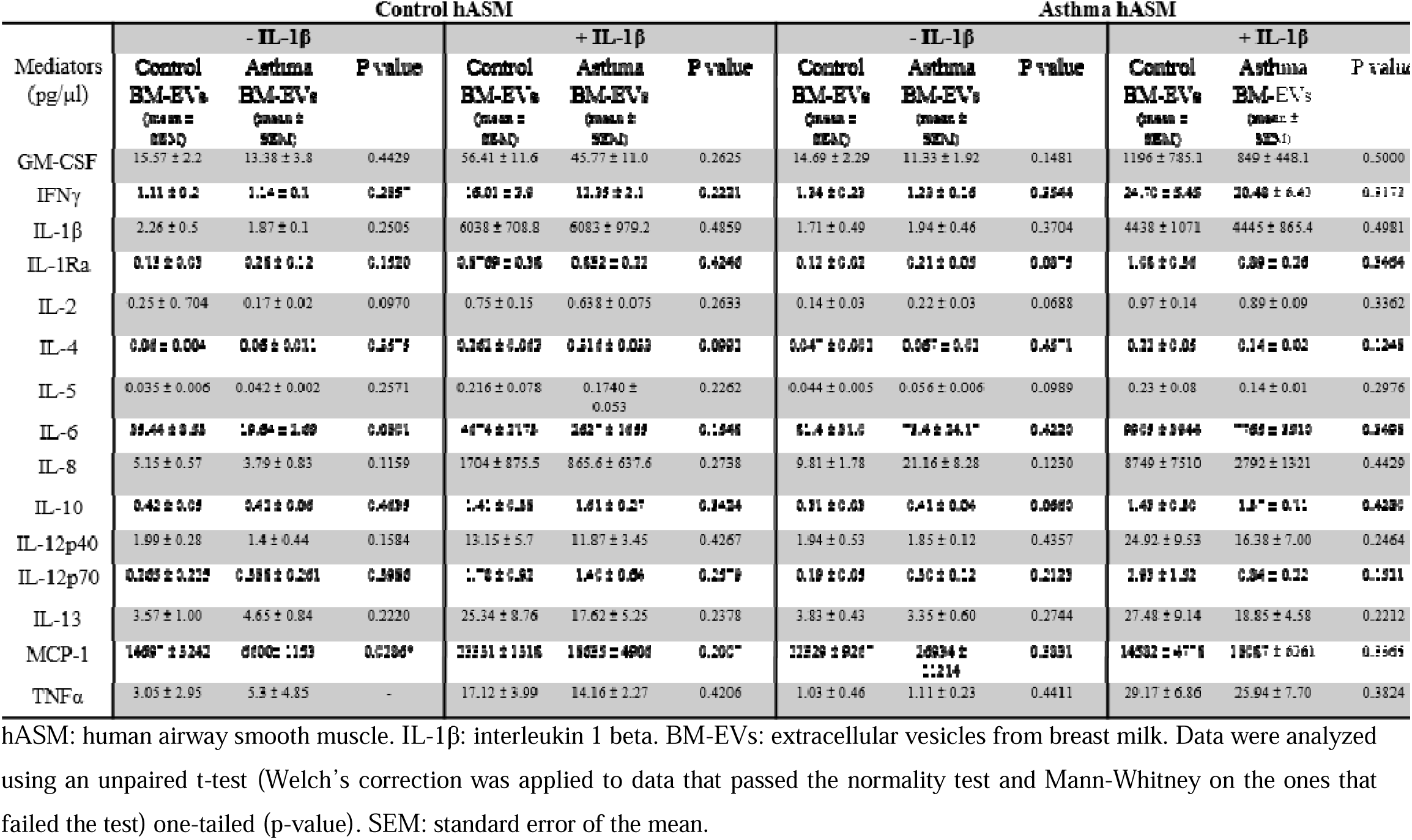
Inflammatory mediators released by control and asthma hASM cells after 48 hours of treatment with control or asthma BM-EVs with or without IL-1 β induction.

## Discussion

Our aim in this study was to isolate and characterize human BM-EVs from mothers with or without asthma, and to investigate whether BM-EVs can potentiate anti-inflammatory effects by regulation of cytokine release from primary hASM cells. Our findings show that average EV size was lower, but stability (zeta potential) was unchanged in BM-EVs from mothers with asthma *vs*. control. Concomitantly, there was an increased concentration of asthmatic BM-EVs, particularly small EVs (<200 nm). Somewhat counterintuitively, asthmatic BM-EVs had lower expression of small EV-specific markers (CD63, CD9, HSP70 and Flotillin-1) than control BM-EVs. BM-EVs exerted differential effects on cytokine release from hASMs in a BM-donor- and recipient cell-specific manner. Treatment with asthmatic BM-EVs reduced pro-inflammatory cytokines (MCP-1, IL-6 and IL-2) in control hASMs, and increased anti-inflammatory cytokines (IL-10, IL-1Ra) in asthmatic hASMs. BM-EVs did not affect cell viability of hASM cells, and both control and asthmatic BM-EVs were taken up hASM cells successfully.

Admyre et al. first identified small EVs in colostrum and mature human milk using electron microscopy and reported the vesicles displayed a typical exosome-like size and morphology of 50 nm (Admyre et al., 2007). Kupusco et al. reported EVs from human breast milk are ∼ 163.5 nm in diameter, with a size distribution of 50-650 nm, and a concentration of 1.5E11 particles/ml (Kupsco et al., 2021). Others have shown EVs consistent with the size (30-300 nm) and morphology of small EVs in human breast milk (Mirza et al., 2019). In our current study, isolated BM-EVs had a size distribution of ∼100-350 nm, with the average size of BM-EVs from asthmatic mothers (135 nm) comparable to healthy mothers (148 nm). TEM imaging revealed a distinct outer membrane and intact spherical-shaped morphology of BM-EVs displaying an approximate exosome size (100 nm). The concentration of BM-EVs in samples from control mothers without asthma (5.13E10 particles/ml) is consistent with previous reports. Intriguingly, we found that mothers with asthma released 4.5 times (2.39E11 particles/ml) more EVs than healthy mothers. We do not know if this is due to an increased biosynthesis and/or enhanced secretion of EVs from cells of origin, nor whether these small EVs are of endocytic or exocytic origin. Zeta potential is a distinguished physical property that indicates the stability of colloidal nanoparticles such as EVs through the measurement of the surface charge (Beit-Yannai et al., 2018; Vogel et al., 2017). We found that zeta potential of BM-EVs presented no difference between mothers with or without asthma (−12mV) and the values were similar to other studies using cow milk-derived EVs (−8.1mV and −17mV) (Carobolante et al., 2020; Shandilya et al., 2020). It is important to note that the better resolution of TRPS *vs*. more traditional approaches (e.g. nanoparticle tracking analysis or NTA) allows precise information about particle size distribution and EV subpopulations (Caputo et al., 2021), and could explain any variations in our data compared to previous work.

The amount of protein yield in each fraction obtained by SEC (F1-12) of BM-EVs increased incrementally across twelve fractions, but remained unchanged between control and asthmatic BM-EVs. To confirm the purity of fractions Western blotting was used to immunoblot for subtype-specific markers for EVs. Only F7-9 of BM-EVs showed a solid abundance of transmembrane proteins enriched in small EVs such as the tetraspanins CD81, CD9 and CD63, and cytosolic proteins (TSG101, flotillin-1, HSP70). Lastly, F9 had lower expression of all these proteins compared to F7-8, probably due to increased soluble proteins and decreased EV concentration. These findings are in concordance with previous work that has reported enrichment of tetraspanins CD9, CD63 and CD81 in human milk exosomes, purified using ultracentrifugation (Admyre et al., 2007; Lässer et al., 2011). In our hands, F1-6 did not express any EV or non-EV markers, as expected and informed by the manufacturer’s instructions. Finally, as described previously, F10-12 presented a high content of protein compared to F1-9, which is attributed to the presence of soluble proteins rather than small EVs, and increased expression of microvesicle-markers MMP2 and ARF6. ARF6 regulates endosomal trafficking acting as a selective recruiter of proteins to the cell surface in microvesicle formation (Gurunathan et al., 2021; Van Niel et al., 2018), while MMP-2 is an important player in EV transport (Das et al., 2021; Shimoda & Khokha, 2017). To assess purity of EV preparations, lipoprotein ApoA1 was used to determine the presence of non-EV proteins co-isolated with vesicle purifications. Fractions 7-9 showed the presence of Apo-A1, but the strongest expression was detected in F10-12. Altogether, we selected F7-9 as a small EV-enriched fraction and used it in all subsequent analysis in the study.

Milk EVs can originate from a variety of cells residing in the mammary gland, such as mammary epithelial cells, immune cells, stem cells, adipocytes and bacteria (Sanwlani et al., 2020). Milk EVs are particularly enriched with proteins associated with trafficking, fusion, and membrane proteins with immunologic implications such as CD9, CD63 and CD81.

While we cannot report which cells our BM-EVs originated from, we note that asthmatic mothers presented with significantly reduced expression of CD63 and flotillin-1 in BM-EVs compared to healthy mothers. CD63 can influence exosome biogenesis, cargo selection, targeting, and uptake, while flotillin-1 is one component of membrane transport and fusion proteins involved in exosome secretion and uptake (Gurung et al., 2021). Whether the differential expression of these proteins impact BM-EV uptake in vivo remains to be elucidated.

To investigate the functional effect of BM-EVs on cellular indices of asthma progression (i.e. inflammation), we performed functional assays to evaluate the physiological effect of BM-EVs on primary hASM cells from donors with or without asthma. hASM cells are considered to play a central role in the pathogenesis of asthma, regulating bronchomotor tone and orchestrating airway inflammation and hyperresponsiveness (Camoretti-Mercado & Lockey, 2021; Tliba et al., 2008). Airway inflammation stimulates the proliferation of ASM cells contributing to the development of hypertrophy and hyperresponsiveness, which is believed to be due to an up-regulation of inflammatory mediators in the airway (Khan, 2013). These cells can release a plethora of inflammatory cytokines, chemokines, and grown factors and express adhesion molecules that are important in modulating submucosal airway inflammation (Chung, 2000; Panettieri, 2002). Conveniently, primary hASM cell cultures have provided an opportune model for studying airway responses in asthma (Johnson et al., 2001; Patel et al., 2012; Sutcliffe et al., 2009; Watson et al., 1998; Wuyts et al., 2003). We observed that BM-EVs exerted differential immunomodulation effects on cytokine release in a BM-donor and recipient cell-specific manner.

Briefly, asthmatic BM-EVs co-cultured with hASMs from control donors decreased pro-inflammatory cytokine secretion: MCP-1, IL-6 and IL-2 *vs.* control-BM-EVs. MCP-1 is an important chemokine released from ASM cells in a culture medium, even under unstimulated conditions. It is chemotactic for monocytes and T lymphocytes, which contribute to airway inflammation in asthma (Sutcliffe et al., 2009). IL-2 has been shown to increase allergic airway responses (Nag et al., 2002), and IL-6 is a pleiotropic cytokine known to have a pro-inflammatory effect in asthma, found in the sputum of asthmatic patients at increased levels after mast cell activation (Shan et al., 2010). Conversely, asthmatic BM-EVs co-cultured with hASMs from asthmatic donors increased secretion of anti-inflammatory cytokines IL-10 and IL-1Ra, pro-inflammatory cytokine IL-2 *vs.* control-BM-EVs. IL-10 is a potent inhibitor of macrophage function and acts by reducing the secretion of pro-inflammatory cytokines such as TNF-α, IL-1, IL-6, IL-8 and MIP-1, and up-regulating the monocyte expression of IL-1Ra, another anti-inflammatory cytokine. In addition, it inhibits the stimulated release of RANTES from human ASM cells in culture, which is a potent chemoattractant for eosinophils, lymphocytes, and monocytes (Chung, 2000; John et al., 1997). Elevated levels of IL-10 have been considered an advantage of steroid therapy and allergen-specific immunotherapy and a target for further studies to improve the treatment of airway inflammatory diseases such as asthma (Böhm et al., 2015; Ogawa et al., 2008). It is important to note that while IL-2 is largely a pro-inflammatory cytokine, it also has anti-inflammatory protective effects in ASM (Hakonarson et al., 1999). We also evaluated the effect of BM-EV treatment in hASMs activated with IL-1β to trigger the release of inflammatory mediators and simulate airway inflammation during asthma. IL-1β is a pro-inflammatory cytokine used to stimulate MCP-1 and other chemokines in hASM cell culture (Wuyts et al., 2003). BM-EV treatments did not exert any effects on cells stimulated with IL-1β, except for an increase in IL-4 secretion observed in control hASMs treated with asthmatic BM-EVs. IL-4 is a T helper type 2 (Th2) associated pro-inflammatory cytokine that induces ASM cells to proliferate and release more inflammatory cytokines, contributing to eosinophilic airway inflammation in asthmatic individuals (Chetty et al., 2021). In summary, BM-EVs modulate the release of inflammatory cytokines in a donor- and recipient-cell specific manner. Whether these trends will be observed in other cell types with BM-EV treatment and if they translate into in vivo adaptations remains to be elucidated.

Our study has some limitations. First, the breast milk samples used in this study are from a longitudinal CHILD Cohort Study, which had been frozen at −80 °C for an extended period prior to the EV isolation and analysis. Second, it is important to note that the milk fat globules were not removed prior to storage, and casein was not precipitated. Casein micelle removal is a common challenge for milk-derived EVs isolation, since they overlap in size with EVs (Welsh et al., 2024). This may have affected the EV population obtained in this study, as milk cell types, casein, and cream-forming milk fat globules can lead to contamination of the natural milk EV population (Zonneveld et al., 2014). In this study, only the fat globule layer and cell debris from milk-thawed samples were removed by centrifugation and filtration processes prior to the EV isolation. Third, we did not have access to colostrum samples to compare the EV composition of milk 3-4 months post-partum as used in this study. Colostrum has a rich composition of immunoglobulins and growth factors that protect newborns against allergic and chronic diseases (Bardanzellu et al., 2017). It is also enriched with small EVs, despite mature milk being an abundant source of intact EVs throughout the lactation curve with no difference between 1-9 months post-partum (Santoro et al., 2023). Thus, colostrum-derived EVs may play an important role in inflammatory modulation. Fourth, the sample size in our study was relatively small, leading to notable variation between biological samples. We highlighted salient findings in the absence of statistical significance, only when the magnitude of the effect underscored its physiological relevance. Future work with a greater sample size may yield statistically significant results. Finally, despite the crucial importance of ASM cells in airway inflammation, other cell types present in the lung such as different types of immune cells, and bronchial epithelial cells, can also participate in asthmatic and allergic processes. In this work, we evaluated the response of human ASM cells to BM-EVs, but it is also important to investigate the biological response of airway epithelium cells and immune cells to these EVs as well.

Overall, our work suggests that the asthma status of the mothers plays an important role in the composition and anti-inflammatory functional activity of breastmilk EVs. BM-EVs derived from asthmatic mothers modulated the inflammatory mediators released by hASM cells in a recipient cell-specific manner. Further investigation is necessary to identify the BM-EVs cargo, and mechanisms underlying the anti-inflammatory effects observed.

## Authors contribution

T.S., T.M.P., S.S., B.B. and P.O.O. performed experiments in the current study, analyzed data, and created figures. T.S., T.M.P. and A.S. helped draft and revise the manuscript. J.G., S.T., E.S., P.M., T.M., P.S., S.R., A.H., and M.A. provided technical and theoretical expertise to complete the work. All authors were involved in manuscript revisions. A.S. designed the project and helped synthesize data, create figures and write and edit the manuscript and its revisions. A.S. is the corresponding author and directly supervised the project. All authors have read and agreed to the published version of the manuscript.

## Acknowledgements

We would like to thank Brian E. Conn (Izon Science, US) for technical assistance with EV biophysical characterization. The authors also like to express our gratitude to our colleagues at the University of Manitoba: Dr. Neeloffer Mookherjee, Dr. Mahadevappa Hemshekhar, Kelsey Fehr and Shana Kahnamoui for their assistance during the study.

## Supplemental material

**Figure S1.**
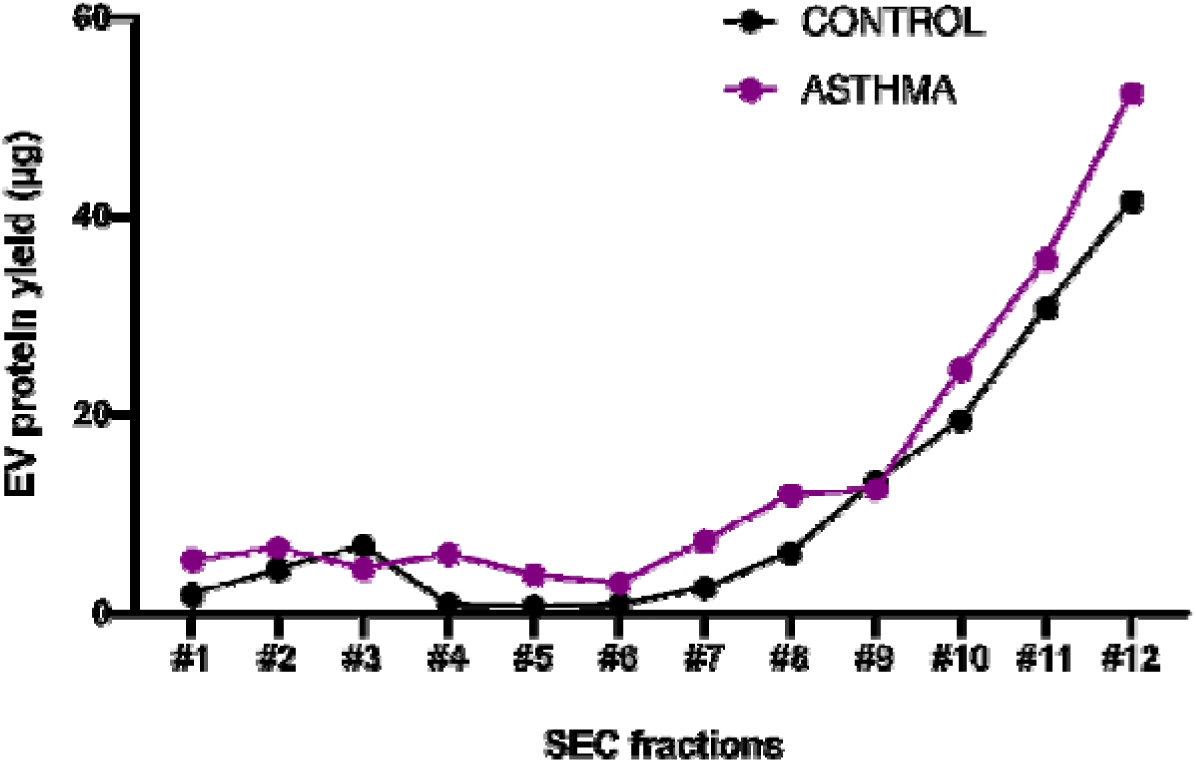
Protein yield (μg) for each SEC fractions (1-12). The protein yield showed an exponential increase in protein concentration from fraction 7 (F7) onwards in both groups.

**Figure S2.**
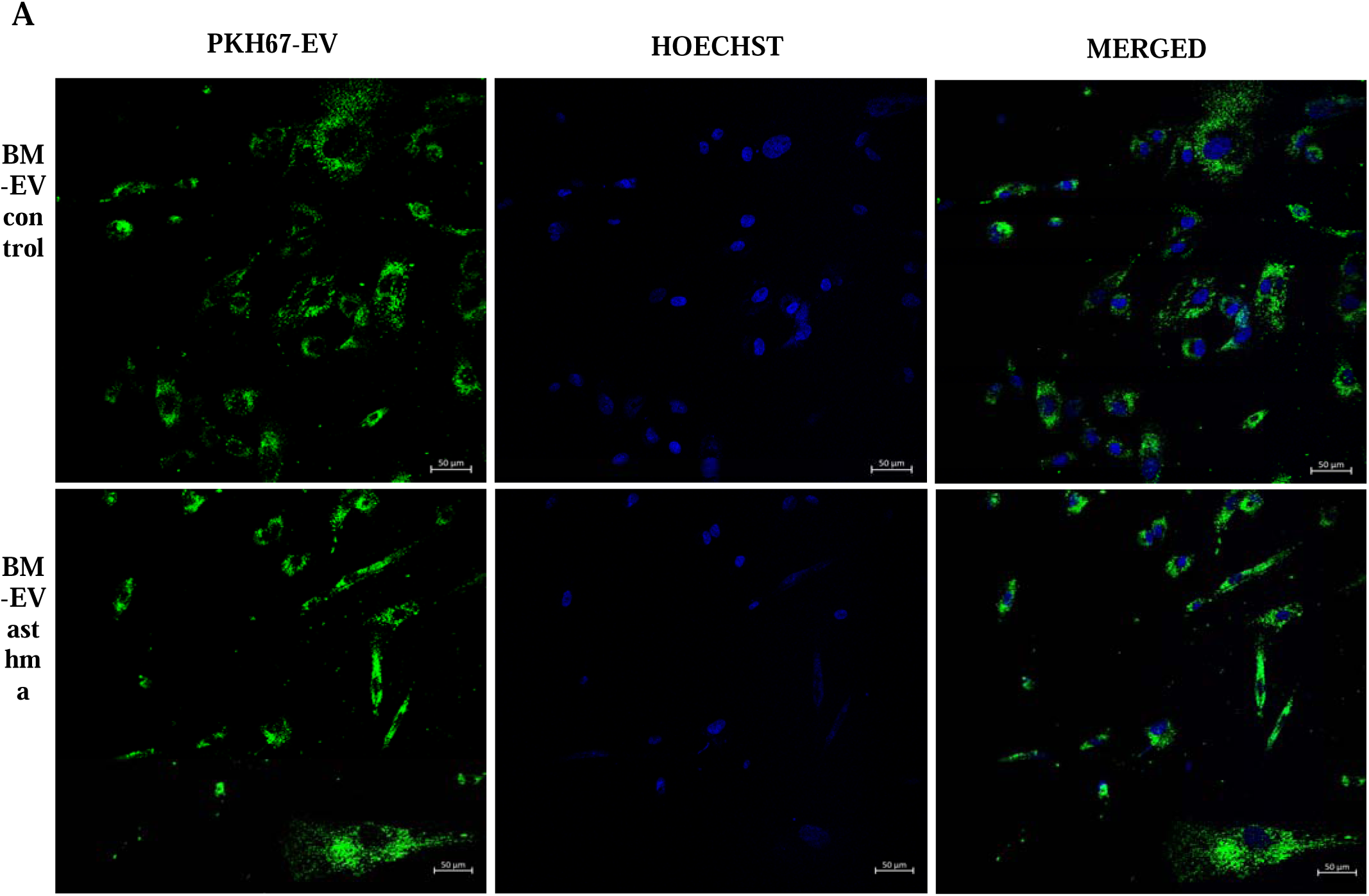
Uptake of BM-EVs labelled with PKH67 by hASM cells. Representative fluorescent images of PKH67 dye (1 µM)-labelled BM-EVs from mothers with and without asthma (control) internalized by hASM cells (green). hASM cells nuclei were stained with Hoechst (blue). Scale bar = 50 μm.

